# Sexual antagonism drives the displacement of polymorphism across gene regulatory cascades

**DOI:** 10.1101/454959

**Authors:** Mark Hill, Max Reuter, Alexander J. Stewart

## Abstract

Males and females have different reproductive roles and are often subject to contrasting selection pressures. This sexual antagonism can lead, at a given locus, to different alleles being favoured in each sex and, consequently, to genetic variation being maintained in a population. Although the presence of antagonistic polymorphisms has been documented across a range of species, their evolutionary dynamics remain poorly understood. Here we study antagonistic selection on gene expression, which is fundamental to sexual dimorphism, via the evolution of regulatory binding sites. We show that for sites longer than 1 nucleotide, polymorphism is maintained only when intermediate expression levels are deleterious to both sexes. We then show that, in a regulatory cascade, polymorphism tends to become displaced over evolutionary time from the target of antagonistic selection to upstream regulators. Our results have consequences for understanding the evolution of sexual dimorphism, and provide specific empirical predictions for the regulatory architecture of genes under antagonistic selection

## Introduction

Adaptive responses to divergent selection in males and females are hampered by a largely shared genome, which slows or even prevents the evolution of sexual dimorphism, where the two sexes reach their respective phenotypic optima. In this situation populations can experience the invasion of “sexually antagonistic”(SA) alleles that are beneficial in one sex, but deleterious in the other^1,2,3,4^. Sexual antagonism is increasingly recognised as a taxonomically widespread and evolutionarily important phenomenon. A wealth of empirical evidence for SA fitness variation across a wide range of animal and plant species has now accumulated^5,6,7,8,9,10,11^ and the balancing selection generated by sexual antagonism in these species helps to maintain surprisingly large amounts of heritable fitness variation^12^. Furthermore, antagonism is thought to be a key driver for the evolution of sex chromosomes^13,14^ and sex determination^15,16,17^, to play a role in reproductive evolution (by eroding “good gene” benefits of sexual selection^18^), and to mitigate the evolution of reproductive conflict between the sexes^19^.

The conditions that favor emergence and maintenance of antagonistic variation in a population have been explored by a large body of theoretical work. These studies have captured the fate of antagonistic variation in infinite populations^20,1^ under a wide range of genetic effects^21^, as well as under selection on linked antagonistic polymorphisms^12,22^, in the presence of genetic drift in finite populations^23,24^ and under fluctuating environments^25^. What they all have in common, however, is that they consider small numbers of allelic variants at one or a small number of loci.

It is important to realise however that the abstract concept of the ’locus’ in these models imposes limitations on the applicability and generality of their results. Specifically, the notion of alleles segregating at isolated and distinct loci makes the implicit assumption that variants with antagonistic fitness effects can arise by simple, individual mutation events. This is appropriate when considering antagonistic selection on protein coding sequences, where non-synonymous substitutions can generate evolutionary relevant phenotypic variation in males and females. However, the assumption of isolated polymorphisms breaks down in the case of regulatory evolution, where the phenotype—and hence fitness—is determined by the match between the sequence of a putative binding site and the motif that is recognised by a transcription factor. Accordingly, it is the combination of sequence states at all positions of a binding site that matters, rather than the state at any individual position. Existing population genetic models of antagonism cannot readily capture this complexity and are thus of limited use to predict antagonistic evolution of gene regulation.

This matters, because antagonistic selection on regulatory regions is all but inevitable. Sexual dimorphism requires the differential use, and hence expression, of genes in males and females and therefore can only arise via a period of opposing selection pressure on the regulation of individual genes. Understanding how dimorphic regulation can evolve, and the antagonistic constraints that may oppose its evolution, necessitates models that can adequately describe the evolution of regulatory binding sites under sex-specific selection. To model binding site evolution, we build on previous work that considers the fitness landscape of sequence states across the entire binding site by integrating the known biophysical properties of TF binding into models of regulatory evolution^26,27,28,29,30,31^.

We extend these models to study the effects of SA selection on cis-regulation. We explore, via simulation and analysis, the selective conditions that permit invasion and maintenance of antagonistic binding site variants in a population. We then expand our modeling framework to consider regulatory cascades under SA selection, and determine where in a regulatory chain polymorphisms are most likely to arise and persist. We show that regulatory architecture has a fundamental impact on our expectations about the selective conditions, and the positions within a regulatory network, that give rise to antagonistic polymorphisms. We further show that antagonistic selection can lead to ongoing reorganisations in regulatory cascades over evolutionary timescales, including abrupt “displacement” events, where the location of polymorphism shifts from genes directly under antagonistic selection, to one of their upstream regulators.

## Results

### A regulatory binding site under SA selection

Gene expression is controlled, to a large extent, by transcription regulation, where transcription factors (TFs) bind to characteristic sequences of DNA (binding sites) upstream of a transcription start site. TFs up- or down-regulate gene expression, for example by aiding or hindering the acquisition of RNA-polymerase at the transcription start site. The biophysical properties of TF binding are well understood^26,27,28,29,30,31^—for a binding site of *n* nucleotides, the relationship between i) the expression level, *E*, of a regulated gene, ii) the number of nucleotides, *k*, in a binding site matched to the maximum binding affinity “consensus sequence”, and iii) the number of TF proteins *P* available to bind to the site is well approximated by Eq. 1 (see Methods).

A gene whose expression is under sexually antagonistic selection experiences conflicting sex-specific pressures on its regulation. We focus on the straightforward case of a somatic gene whose expression is selected to be maximum in males and minimum in females (the sign associated with the selection pressures operating on each sex is arbitrary and identical results would be obtained for the opposite case). We begin by focusing on a single binding site that up-regulates the expression of its target, meaning that high affinity binding sites are favored in males and low affinity sites in females (Figure 1). Eq. 1 thus provides us with the basis for an empirically grounded genotype-phenotype map for this system, since it relates the nucleotide sequence at the binding site to the expression level of the gene under antagonistic selection. We assume that the level of gene expression *E* relates to fitness by a sigmoidal function (see Methods, Eq. 2) which increases from 1 − *s_m_* (when *E* = 0) to 1 (when *E* = 1) in males and decreases from 1 (when *E* = 0) to 1 − *s_f_* (when *E* = 1) in females.

**Figure 1:**
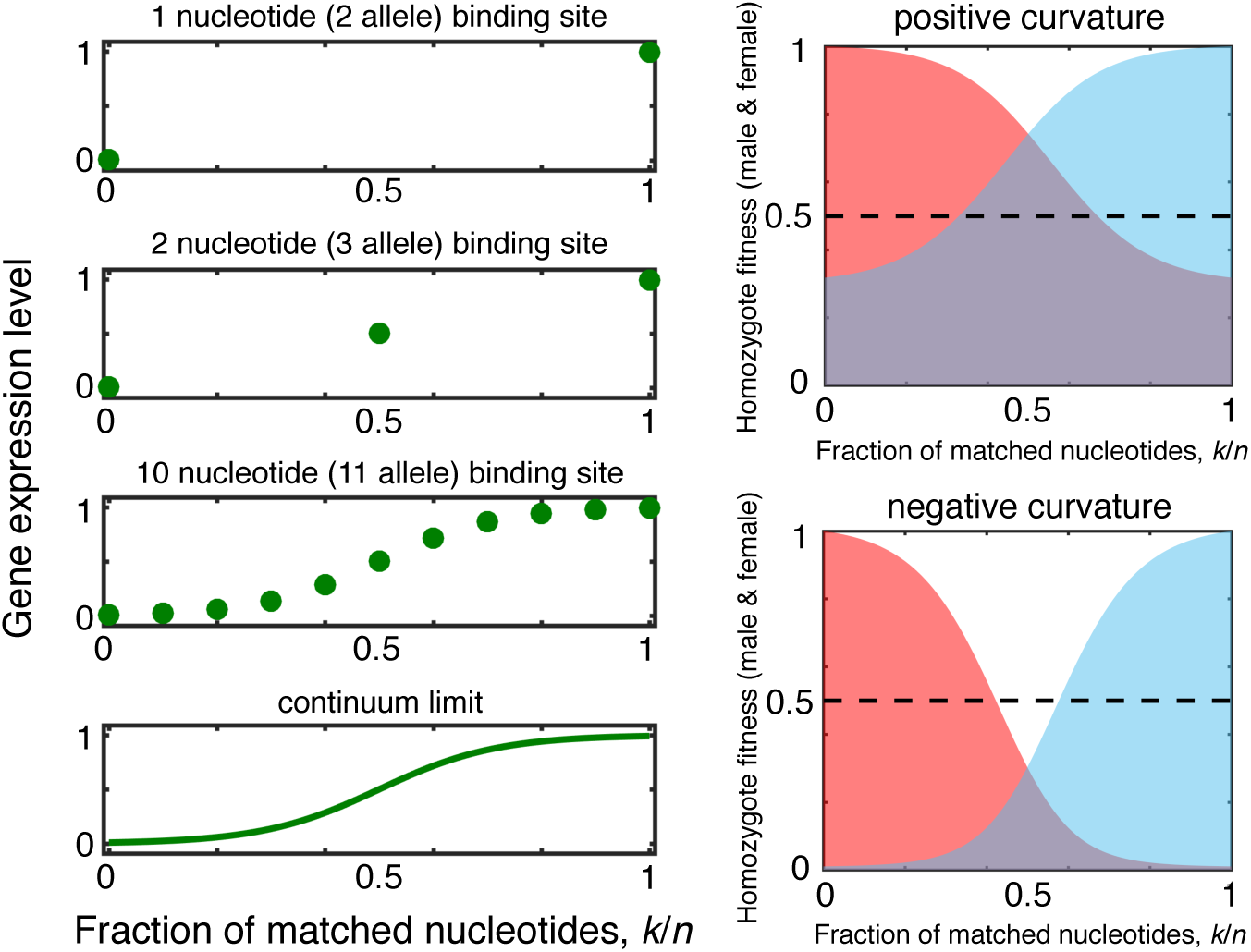
Sexually antagonistic selection on gene expression. Regulation of gene expression by TF binding sites is well understood at a mechanistic level, allowing us to construct explicit genotype-phenotype maps. In the case we consider, expression level *E* increases with the number of nucleotides *k* correctly matched to a consensus sequence (left). Binding site length *n* is well known to have important consequences for the dynamics of binding site evolution^31,30^ generally. However, population genetic models of sexual antagonism typically focus either on a 2-allele system^20,1,21,23,24^ (corresponding to a binding site of 1nt in length), or in some contexts on the continuum limit and infinite alleles^32^. Eukaryotic TF binding sites, in contrast, are typically around 10 nucleotides long^30^, and vary from as short as 5nt to > 20nt in some cases. By varying the binding site length *n* we can generate a system with as few as 2-alleles (top - left) to an infinite number of alleles in the continuum limit (bottom - left). Real eukarytoic TF binding sites of *n* = 10nt results in 11 alleles at a given locus. We assume that expression is selected to be high in males (blue) and low in females (red) (right), and we consider fitness landscapes with different “curvatures” corresponding to different levels of average fitness at intermediate expression levels.

The relative steepness of male and female fitness functions has important consequences for the evolutionary dynamics of antagonistic binding site variants. In particular, we must distinguish SA fitness landscapes with *positive* and *negative* curvature, where curvature is determined by average fitness at intermediate expression levels (see Methods). Curvature is said to be positive when the average fitness across males and females of intermediate expression alleles (*E* = 1/2) is greater than the average fitness of maximum or minimum expression alleles (*E* = 1 or *E* = 0) and to be negative when the converse is true (Fig. 1 - right).

We begin by determining, via individual-based simulations (see Methods), the stationary distribution of binding sites, and the resulting expression polymorphism, in antagonistic fitness landscapes with both positive and negative curvature as a function of binding site length *n* and population size *N* (Figure 1).

We explore the action of antagonistic selection on gene expression for binding sites across a wide range of lengths 1 ≤ *n* ≤ 100 (Fig. 2 - left). We find that high levels of expression polymorphism always evolve in landscapes with negative curvature (i.e., when intermediate expression is deleterious on average compared to high or low expression). Conversely, high levels of polymorphism never evolve in landscapes with positive curvature (i.e., when intermediate expression is on average fitter compared to high or low expression), with the notable exception of the limiting 2-allele case (*n* = 1nt). These results hold over a wide range of population sizes 10^2^ ≤ *N* ≤ 10^4^ (Fig. 2 - right).

**Figure 2:**
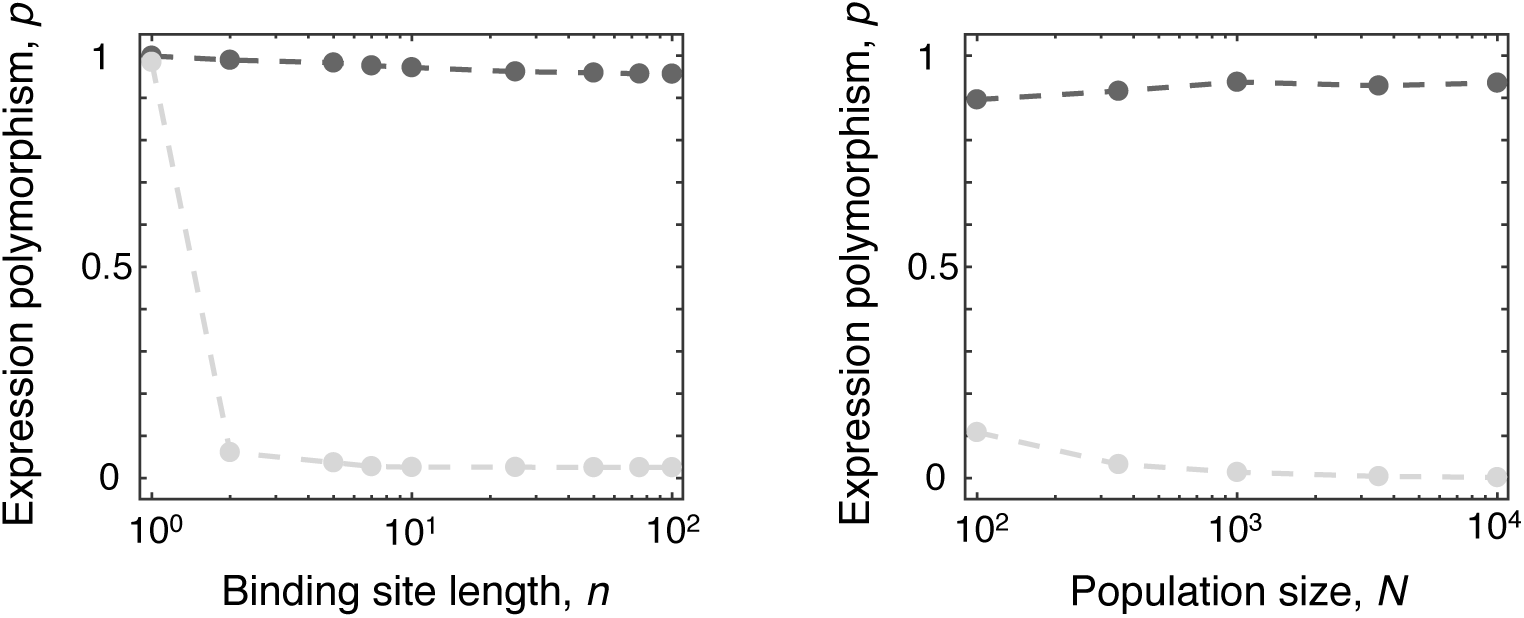
Expression Polymorphism at a single binding site. Results of individual based simulations showing the amount of polymorphism (*p*) as a function of (left) binding site length (*n*) and (right) population size (*N*) for landscapes with negative (dark gray) and positive (light gray) curvature. Points show the ensemble average of 10^4^ runs at each parameter value. Default population size was fixed at *N* = 10^3^ and default binding site length at *n* = 10. Per-binding site mutation rates were set to 4*N µ* = 0.1, selection was assumed to be strong (*s_m_* = *s_f_* = 0. 1). Curvature was set to *c* = ±0.2 and the fitness landscape had steepness *h* = 10 (see Methods). Each simulation was run until 10^6^ mutations per binding site had occurred.

Figure 3 illustrates the intuitive explanation for the effect of fitness landscape curvature, in terms of the selection gradient experienced by mutations that increase or decrease binding affinity in a typical binding site of 10nt. When curvature is negative, polymorphism is favored between pairs of alleles with intermediate binding strength, *and* the polymorphisms are subject to divergent selection gradients, with weaker sites favored to get weaker and stronger sites to get stronger. This results in disruptive selection which generally leads to polymorphism. When curvature is positive, polymorphism can still sometimes be favored at intermediate expression levels, but there is no disruptive selection and alleles of intermediate binding strength are maintained. This is because SA can be resolved to the mutual advantage of both sexes by fixing an allele of intermediate expression that maximises average fitness across males and females. In the 2-allele case, landscape curvature does not result in these contrasting dynamics. This is because when *n* =1 binding is a binary function of whether or not the binding site matches the consensus sequence, meaning intermediate binding is not possible. Thus, in this case—even with positive curvature—polymorphism is maintained.

**Figure 3:**
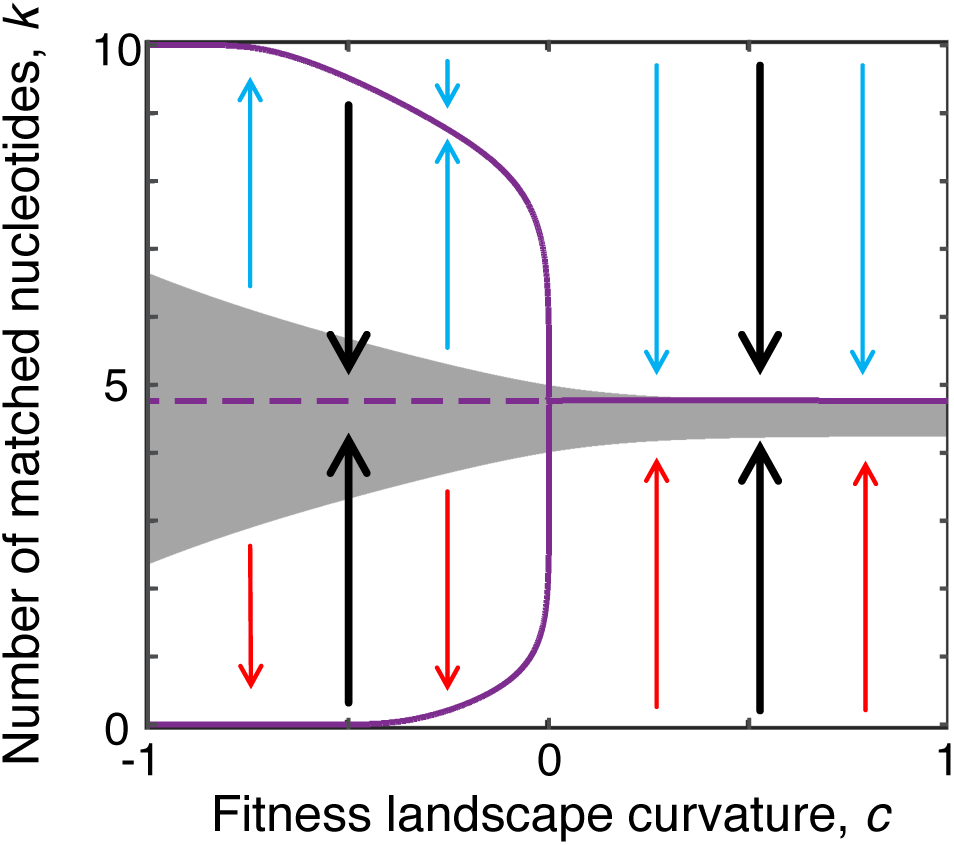
Pairwise invasion plot for a single binding site. We calculated the selection gradient (see SI) for a “typical” binding site of 10 nucleotides, assuming weak mutation so that at most two alleles segregate in a population at a given time, as a function of fitness landscape curvature *c*. We also used a two-allele approximation to determine whether polymorphism was favored (see SI), with the polymorphic region indicated in dark gray. Solid purple lines indicate stable monomorphic equilibria in the dynamics under uni-directional selection while dashed lines indicate unstable equilibria. Blue arrows indicate the direction of the selection gradient on males and red the direction of selection on females. Black arrows indicate the direction of evolution in a monophonic population under antagonistic selection (see SI). When curvature is negative (left hand side, *c* < 0) there are two distinct stable monomophic equilibria corresponding to high fitness males and low fitness females (*k* → *n*) or low fitness males and high fitness females (*k* → 0), a scenario that results in the maintenance of SA polymorphism. When curvature is positive (right hand side *c* > 0) there is a single intermediate equilibrium corresponding to males and females with high fitness i.e. antagonism is resolved.

### Polymorphic displacement in a regulatory cascade

We have focused so far on a single binding site at a single target gene. However, most genes are regulated by multiple binding sites and most regulators are themselves subject to regulation, as part of a wider regulatory network^33,34^. This is particularly true for genes involved in sex determination and sexual differentiation for example, which are frequently arranged in regulatory cascades^35^. In relation to antagonistic selection, this regulatory connectivity creates the potential for polymorphism to arise at multiple points in a regulatory cascade, even if only a single downstream gene is subject to direct SA selection for expression.

In order to investigate the invasion and maintenance of antagonistic polymorphism across regulatory cascades, we once again assume a gene whose expression is under antagonistic selection such that high expression is favored in males and low expression in females. However we now assume that this gene (gene 1) is at the bottom of a three-gene regulatory cascade (Figure 4c, right), where its expression is up-regulated by a second (gene 2) which in turn is up-regulated by a third (gene 3). The third gene further has a binding site that up-regulates its own expression in response to some constant input signal (see Methods).

**Figure 4:**
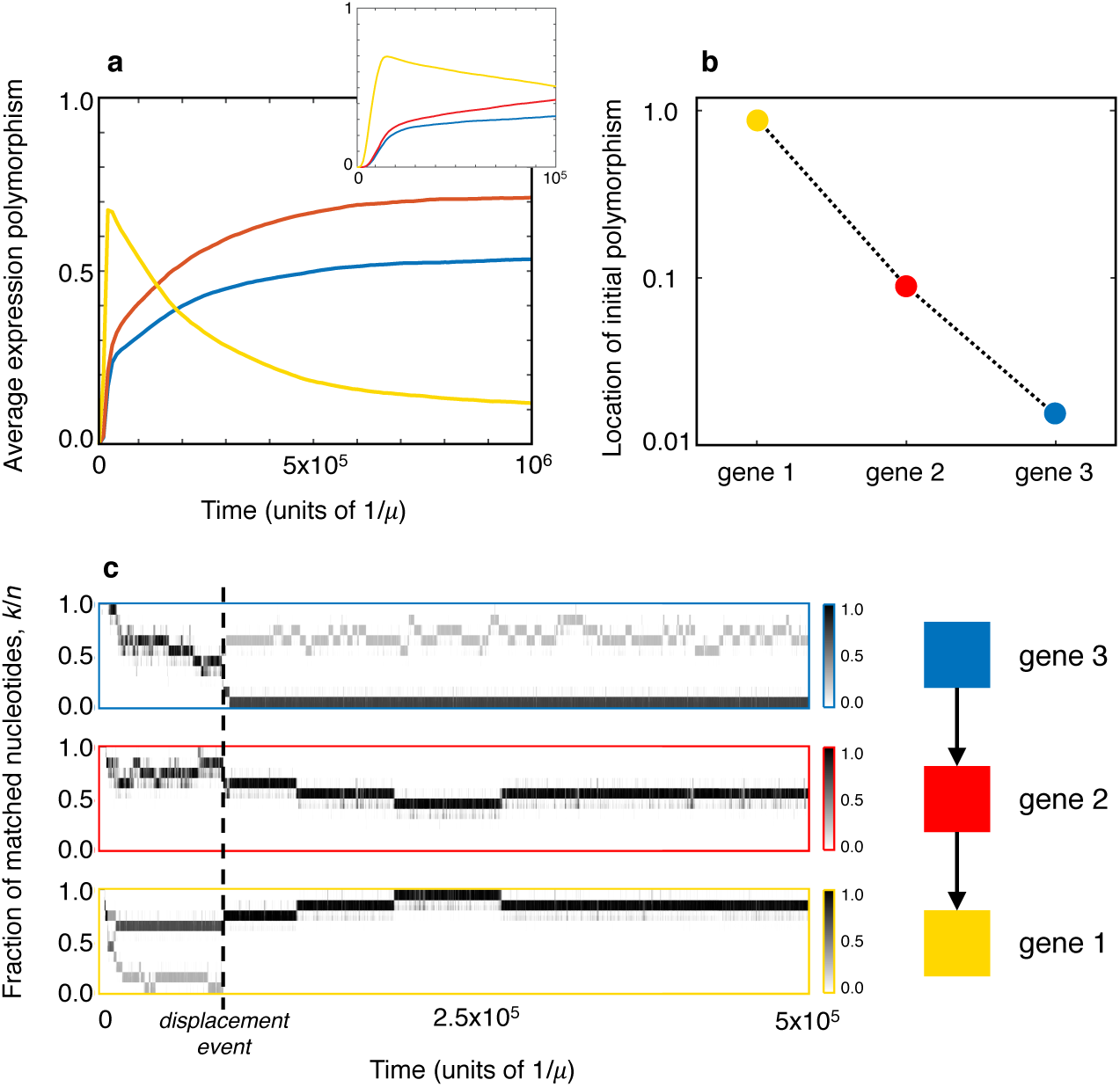
Displaced polymorphism in a regulatory cascade. a) We observed the average expression polymorphism for each gene over evolutionary time. The initial phase (inset) sees polymorphism arise at gene 1 (yellow) and beginning to pass to gene 2 (red) and gene 3 (blue) higher up the cascade. b) We determined where in the cascade polymorphism of *p* > 0.5 first arose. In > 90% of simulation runs polymorphism initially arises at gene 1, with the frequency declining approximately exponentially with position in the cascade. c) A sample path for a single simulation run shows the dynamics of displacement explicitly, with gene 1 quickly acquiring polymorphism until a displacement event shifts the polymorphism up the chain to gene 3. These individual based simulations for a cascade of three genes were carried out using the same default parameters given in Figure 2.

Under a fitness landscape with negative curvature, antagonistic selection at gene 1 could potentially lead to polymorphism at any of the three binding sites in the cascade. However, determining precisely where polymorphism will arise is not straightforward, since there is a great deal of epistasis between mutations at different positions in the cascade, meaning that both the ordering of mutations as well as their average fitness effects in males and females becomes important to subsequent evolutionary dynamics^36^.

Starting with a three-gene cascade in which all binding sites have high affinity (*k* = *n*), we explored the evolutionary dynamics of all three binding sites using simulations (Figure 4a). We observe that a high degree of polymorphism initially arises at gene 1, only to subsequently shift towards genes 2 and 3 that sit higher up the cascade. This non-monotonic behavior of expression polymorphism at gene 1 arises due to a phenomenon of *polymorphic displacement*, in which SA initially generates polymorphism at the gene directly under antagonistic selection, only for the polymorphism to become displaced to another gene in the cascade, which remains polymorphic over longer timescales.

To understand these surprising dynamics we must first look at both the fitness of heterozygotes in the presence of polymorphism and the evolvability of SA variants at different points in the regulatory cascade. We observe (Figure 4b) that polymorphism almost always initially arises at gene 1 (> 90% of cases), and that there is an approximately exponential decline in the frequency of initial polymorphism as we move up the cascade to genes 2 and 3. The reason for this decline lies in the fact that the exact amount of regulatory protein has only a marginal effect on the expression of a downstream gene, as long as the proteins are reasonably abundant and binding is strong. As a consequence of this buffering effect, mutations to the binding sites of genes 2 and 3 (which affects the amount of protein produced from these genes) initially have little effect on the expression of the downstream gene 1 and thus selection gradients on the binding sites at genes 2 and 3 are comparably weak. Accordingly, it is relatively easier for initial polymorphism to arise at gene 1, where the effects of mutations on the selected expression level are strongest (see SI).

Once polymorphism takes hold, however, the overall fitness of polymorphic variants arising at gene 1 is lower than of those arising at gene 2 or gene 3. This is also due to the tendency for genes lower in the cascade to compensate for a change in the expression of genes higher up. If gene 1 has a strongly polymorphic binding site such that homozygotes have either expression *E* = 1 or *E* = 0, the heterozygotes have average expression *E* = 0.5. However, if the equivalent polymorphism arises at gene 2, the homozygous cases still correspond to gene 1 having either expression *E* = 1 or *E* = 0, while the heterozygote typically has expression *E* > 0.5 because, again, the effect of reducing the amount of an upstream regulatory protein is somewhat buffered by the regulation of gene 1. As a consequence of this, the average fitness across males and females of heterozygotes at gene 2 and gene 3 with *E* > 0.5, in the landscape with negative curvature, is greater than that of heterozygotes at gene 1 with *E* = 0.5. Thus, displacement initially arises due to a direct fitness advantage to polymorphisms arising at points higher up in a regulatory cascade (see SI). Finally we note that over long timescales, polymorphism is more likely to ultimately reside at gene 3 than at gene 2. As we show in the SI, this arises because genes higher up a cascade are better able to reduce fitness variation between males and females and thus tend to be more stable.

A typical example of the polymorphic displacement phenomenon is shown in Figure 4c. Here, polymorphism arises quickly at gene 1 before being displaced to gene 3, which remains polymorphic over many generations. As Figures 4a and 4c both illustrate, displacement takes place over long evolutionary timescales, with binding sites experiencing around 10^5^ mutations before any displacement occurs. We are thus describing a slow and ongoing reorganization of regulatory cascades in response to SA selection. We also note that the phenomenon we describe is expected to be a general feature of landscapes with negative curvature (see SI). In this type of landscape, more extreme heterozygote expression levels are advantageous, and tend to arise at genes higher up regulatory cascades.

## Discussion

The regulation of gene expression is not only a prime mechanism by which sex-specific adaptation can be achieved, but also an inevitable target for sexually antagonistic selection. By integrating the population genetics of antagonistic variants with a biophysical model of transcription factor binding, our study has generated a number of important predictions for the dynamics of regulatory evolution under antagonistic selection.

First, we show that for binding sites of realistic length, sexually antagonistic polymorphism will only be maintained when intermediate expression levels are, on average, deleterious compared to high or low expression levels. In this scenario of negative curvature, the fitness landscape generates disruptive selection at intermediate binding that will favor segregating binding site variants of ever more extreme affinities. Importantly, the effect of fitness landscape curvature vanishes for the extreme case of binding site length *n* = 1, the situation captured by standard 2-allele models and antagonism. At this limit, mutational effects on TF binding are so coarse that alleles with intermediate expression cannot arise, and antagonistic polymorphism is predicted even with positive curvature.

Second, our model allows us to gain insight into the distribution of polymorphism across regulatory cascades. We predict that allelic variation will be subject to displacement along the regulatory hierarchy. While polymorphisms are most likely to arise at the target of selection, they can subsequently move to other genes higher up the regulatory cascade. The ultimate location of polymorphism is expected to be that which offers the greatest average fitness to heterozygotes while minimizing fintess variation between the sexes (see SI). In the type of cascade modeled here, this corresponds to the gene at the top of the regulatory chain, where buffering of regulatory effects in heterozygotes results in expression *E* > 0.5 at the target gene and an associated benefit compared to heterozygotes with strong and weak binding alleles at the target gene.

While we predict that polymorphism at the top of a cascade will be most beneficial, it is worth noting that this expectation rests on a number of assumptions that do not always hold in real systems. As our simulations show, the timing of displacements is highly variable and the level at which polymorphism occurs will depend on the time of observation. Furthermore, the displacement of polymorphism is a highly stochastic process. Even when assuming strong selection, the fitness differentials that drive upward displacement rapidly decline along the cascade. Thus, while displacing polymorphism away from the target gene generates significant gains, the location of polymorphisms in the higher echelons of the cascade that we observe in our simulations is largely stochastic and dictated by where suitable mutations first arise. It is likely that this tendency will be exacerbated in real regulatory systems, where the fitness effects of regulatory mutations may have significant pleiotropic effects. As a consequence, it will be difficult to make precise predictions about the location of polymorphism, other than that it tends to be above the downstream target gene.

Perhaps a more significant factor that will impact displacement is the structure of a regulatory network. Our simple linear cascade assumes a single target gene under antagonistic selection, yet real-life regulatory networks may feature multiple targets. In cases where all of these targets are under similar antagonistic selection, such a modular organization may favor and precipitate upward displacement of regulatory polymorphism, because modularity amplifies the selective benefits of upstream regulatory variants whose effects propagate across all downstream target genes. In cases where selection differs between co-regulated targets (some antagonistic, some directional/stabilising, or antagonistic selection of opposing directions), in contrast, altered regulation of upstream TFs may generate deleterious pleiotropic effects and prevent polymorphism from being displaced. We may then either see the persistence of antagonistic polymorphism at individual target genes or larger-scale rewiring of gene regulatory interactions to create modules of genes under similar selection (see e.g.^37^).

The question of where antagonistic polymorphism most likely resides within regulatory networks has consequences for our understanding of how antagonism is resolved—and hence how sex-specific development is regulated. It is widely assumed that over the long run, sexual antagonism is resolved by mechanisms that maintain the benefit to one sex while removing the cost to the other. The evolution of sex-specific regulation in antagonistic genes is a prime mechanism to achieve this, certainly in the case that we consider here, where adaptive conflict between the sexes occurs over expression levels (rather than coding sequence) of a gene. Our prediction of upwards displacement of antagonistic polymorphism therefore also implies that we should expect to find the sex-specific regulation resolving antagonism to occur at higher levels of the regulatory hierarchy. Reflecting the arguments on modularity above, this should particularly be the case where genes under antagonistic selection are organised into co-regulated modules. Not only should upward displacement of polymorphism be more strongly selected in these cases, but also its eventual resolution.

Our work has shown that antagonistic selection acting on gene expression can give rise to counter-intuitive evolutionary dynamics across regulatory networks. These are driven by the conflicting impacts of the inherent robustness of networks, whereby changes to the expression of an upstream regulator are frequently compensated for by others downstream. Such buffering tends to prevent the emergence of initial polymorphism at upstream genes, but once such polymorphism exists at downstream targets, favors its upward displacement. Over time, we would therefore expect both antagonistic polymorphism and the sex-specific regulation that may arise to resolve it to reside in the upper reaches of regulatory networks. Testing these predictions directly is difficult, as current data on antagonistic loci and sex-specific resolution are relatively sparse. Interestingly, however, parallels exist between sex-specific selection pressures and directional selection in fluctuating environments^38^. It is therefore plausible that evolutionary dynamics analogous to those described here occur in networks governing the response to alternating environmental conditions, allowing the use of microbial evolution for experimental tests of our theory.

## Methods

Here we describe the details of the biophysical and population genetic model used to generate our results. Transcription factor binding sites are typically around 10 nucleotides long in eukaryotes^30^, while the per-nucleotide substitution rate in *Drosophila* is 4*N u* ~ 0. 01 and an order of magnitude lower in humans, placing both species in the weak mutation limit. For simplicity in our simulations (which vary population size, binding site length and the number of binding sites) we assume a “standard” binding site of length *n* = 10 and set the per-binding-site mutation rate at *µ* = 10*u*. We then run all of our simulations with 4*N µ* = 0. 1, which keeps all of our simulations in the weak mutation limit^39,40^.

### Gene Expression

The biophysics of transcription factor binding is well approximated by assuming an optimal consensus sequence, such that each nucleotide in a contiguous sequence of *n* nucleotides can be considered as either “matched” to the consensus sequence or not, with a matched nucleotide independently contributing an amount, *ϵ* ~ 1 − 3*k_B_T* ^26,41^, to the site’s binding energy. The probability *π_k_* that a site consisting of *k* nucleotides is bound by a TF protein is given by
@@
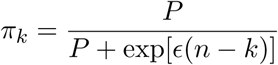

where *P* is the number of TF proteins available to bind to the site. We assume *P* = 200 in our simulations. The rate of transcription (for a fixed decay rate) and the number of translated proteins at the target for a site that up-regulates expression can then be treated, to a first approximation, as proportional to the probability that the site is bound. If we define the expression *E* of the target gene as the number of expressed proteins proportional to the maximum we have simply
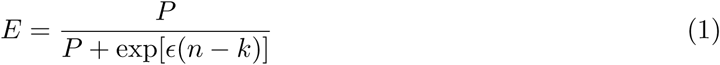

Mutations to binding sites are assumed to occur via single nucleotide substitutions, such that the probability of increasing the number of matches by 1, from *k* → *k* + 1, is *u* (*n* − *k*)/3 where the factor 3 reflects the fact that only one out of the three non-synonymous mutations correctly match to the consensus sequence. Similarly the rate of mutations that decrease the number of nucleotide matches by 1, from *k* → *k* − 1 is *uk*. We therefore not only have a multi-allele system but one with asymmetric forward and back mutations, which makes analytical treatment difficult in most cases.

Since we are considering diploid organisms we in fact have two alleles, 1 and 2, with two expression levels *E* _1_ and *E* _2_, so that overall expression at a locus is given by *E* = 0. 5(*E* _1_ + *E* _2_).

### Fitness Landscape

As described above we assume that the gene under SA selection is favored to have high expression in males and low expression in females, with fitness in both sexes following a sigmoid function of expression levels:
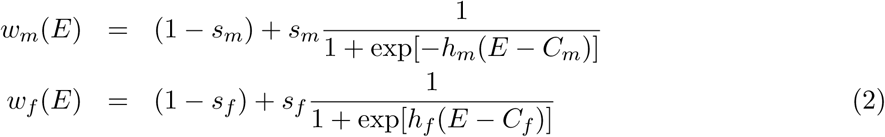

where *w_m_* (*E*) is male fitness and *w_f_* (*E*) is female fitness, *s* defines the overall strength of selection, *h* determines the steepness of the sigmoid function and *C* determines the position of the threshold— where the contribution of expression to fitness is half its maximum. We can then define
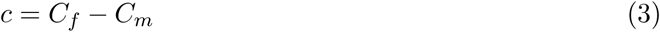

as the curvature of the landscape, so that if *C_f_* > *C_m_* the average effect of an allele with intermediate expression *E* = 0. 5 will be beneficial compared to alleles with high (*E* = 1) or low (*E* = 0) expression.

Below we give further details of the modeling framework, simulations and analytical results used to produce the results presented in the main text. We begin by describing evolution of a binding site under a biophysical model for transcription regulation in the presence of sexually antagonistic selection on a target gene. We then describe the evolutionary dynamics of the model under weak mutation and the conditions that lead to polymorphism between pairs of alleles. These models and analytical approximations are derived for a single (diploid) target gene. Finally, we describe an extension to our model to include multi-gene cascades and the conditions that lead to displaced polymorphism in individual-based simulations under the diploid Moran process.

### Gene regulation under weak mutation

We employ a simplified model of transcription regulation in which transcription factor binding sites are composed of a contiguous region of *n* nucleotides, such that at each position there is a “correct” nucleotide that contributes an amount *ϵ* to the site’s binding energy, while an “incorrect” nucleotide contributes nothing. We assume that *ϵ* is constant across positions, and each position contributes equally and independently to the site’s binding energy. The probability that the site is bound is then given by Eq. 1 in the main text. While in reality the assumption that each nucleotide position contributes equally and independently to binding energy does not necessarily hold^26,41^, this simplified model adequately describes the biophysics of transcription factor binding (and generates a probability of binding that is (including the sigmoidal relationship between binding probability and the number of correct nucleotide matches) and therefore captures the evolutionary dynamics of gene regulation^26,27,28,29,30,31^.

The degeneracy among genotypes assumed by our simplified model allows us to reduce the number of different alleles associated with a given site from 4*^n^*—the number of distinct genotypes that can occur—to *n* + 1—the number of possible “correctly matched” nucleotides at the site. A binding site is thus characterized by a single number *k* which is the number of correctly matched nucleotides. Mutations occur through single nucleotide substitutions which increase, 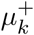, or decrease, 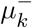, the number of matches at the site from a starting value of *k*.The rates of these mutations are
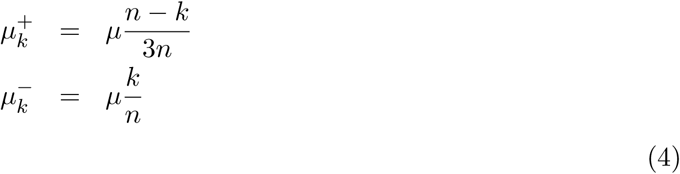

where *µ* is the per-nucleotide rate of substitutions. What is immediately clear from Equation S1 above is that the rates of mutation increasing and decreasing the number of matches are asymmetrical and vary with genotype. This is a violation of the assumptions made in most simple population genetic models (of SA and more generally) and precludes the standard analytical treatment of allelic dynamics using a diffusion approximation (because the resulting system of differential equations cannot be solved). However, because we are considering mutations at the level of nucleotide substitutions with rates that are typically as low as 10^−9^ − 10^−7^ ^42^, we can treat the evolution of a binding site in the weak mutation limit, i.e., in the limit where the product of effective population size and mutation rate is small, or more accurately 2*nµN_e_* ≪ 1.

In the absence of SA, evolutionary dynamics in the weak mutation limit are well approximated if we assume that the population is monomorphic, and that a given mutation has time to either reach fixation or be lost before another arises. Thus, we can simply calculate the fixation probabilities of single mutations while ignoring the effects of clonal interference. If we write 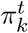 for the probability that the population has a binding site with *k* matched nucleotides at time *t*, then in the weak mutation limit we can write
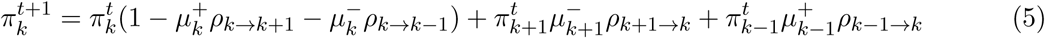

where *ρ_i_*_→*j*_ is the probability that a mutant with *j* matches fixes in a population where a inding site with *i* matches is resident. For a given pair of alleles in the absence of SA, this fixation probability is given by Kimura’s expression^32^. At equilibrium, we can then use detailed balance to find the following recursion relation
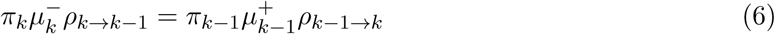

which can be solved numerically.

When we are dealing with SA, this weak mutation treatment unfortunately breaks down. With balancing selection potentially acting on polymorphisms, allelic dynamics are too slow for populations to be assumed to be monomorphic and the evolutionary dynamics of an invading allele among males and females is in general different. We can get around this by making the additional assumption that selection is weak. Polymorphism then cannot typically be maintained for prolonged periods of time and an allele under antagonistic selection is at approximately equal frequencies in males and females^43^. Given this, we can use Equation S3 above to gain insight into the evolutionary dynamics of SA at a binding site under weak selection.

### Mutation-selection gradient

To analyze the evolutionary dynamics of a binding site under in the weak mutation weak selection limit, first recall Kimura’s^32^ expression for the fixation probability of a mutation with relative fitness 1 + *s* against a resident with fitness 1:
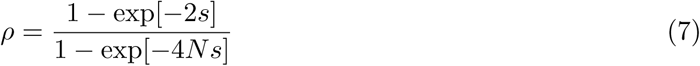

where weak selection requires 2*Ns* ≪ 1. We then use Equation 2 of the main text to calculate the average fitness effect of a mutation that changes the number of nucleotide matches in a binding site by ±1:
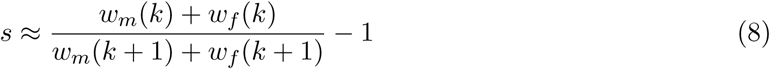

where *k* is the number of nucleotide matches in Equation 1 and *w*(*k*) is the fitness for a homozygote with *k* nucleotide matches. Substituting this into Equation S3 we can calculate the ratio of transition probabilities for mutations that increase or decrease nucleotide matches
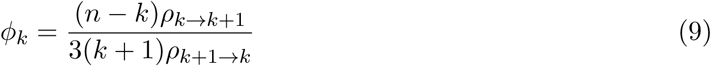

If *ϕ_k_* > 1, the number of nucleotide matches tends to increase whereas when *ϕ_k_* < 1, the number of nucleotide matches tends to decrease. This can be used to describe the direction of evolutionary change of a binding site under uni-directional selection, as shown in Figure 3 of the main text.

### Polymorphism

In order to measure polymorphism in a multi-allele system we calculate the expression difference across alleles at each locus in each individual, which gives us the following expression for polymorphism *p* at a given locus:
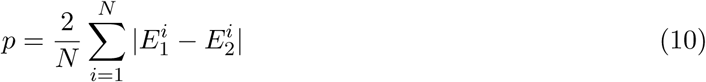

where 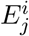 is the expression level of allele j (1 or 2) in individual *i*. If there are two alleles segregating at equal frequency, one with maximum expression and one with minimum expression, this will result in polymorphism *p* = 1 (since approximately half the population will be heterozygous).

We used the results of^43^ to determine whether a given pair of neighboring binding site variants segregating in a populations are favored to be polymorphic. This condition is given in^43^ as
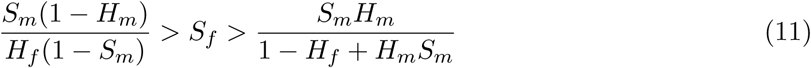

where we have
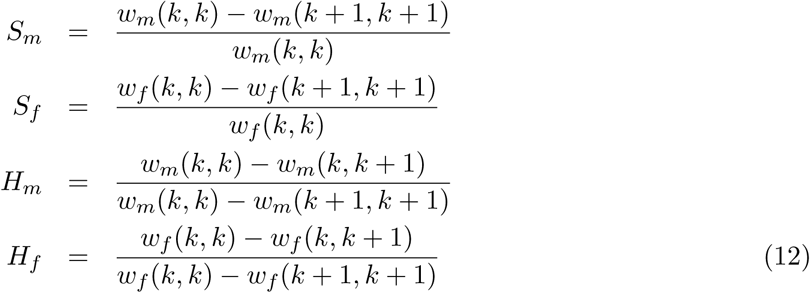

We use Equations S8 and S9 together to determine whether polymorphism will arise in binding sites, as shown in Figure 3.

In contrast to the polymorphic case, the conditions for fixation of a mutant allele in a single locus, two allele system are
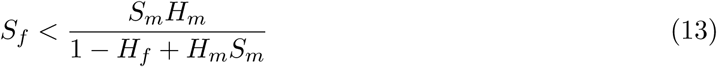

for a male-beneficial mutation and
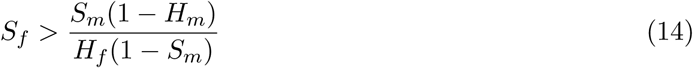

for a female-beneficial allele. This is used to describe the evolutionary dynamics for a male- or female-beneficial binding site variant as shown in Figure 3 of the main text.

### Continuum limit

To understand the evolutionary dynamics of binding sites described in Figure 3 of the main text it is instructive to consider the limit of continuous gene expression and small mutations. In this case Equation 9 above becomes
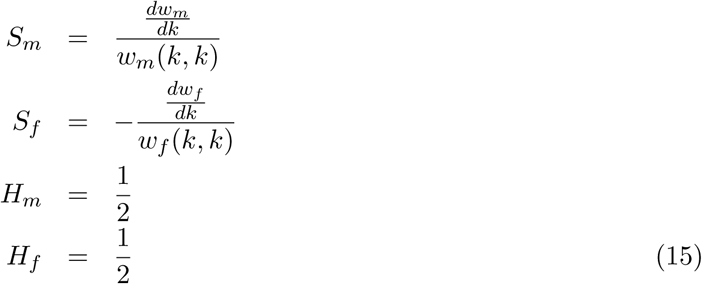

and the condition for invasion of a male beneficial allele becomes
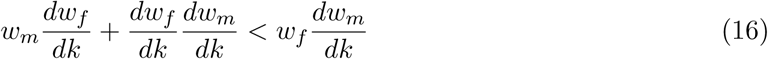

If we then take Equation 2 of the main text under the limit of strong selection, symmetrical selection, i.e. *h_m_* = *h_f_*, *s_m_* = *s_f_* = 1 we recover 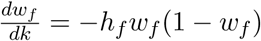 and 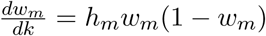 to give
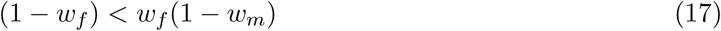

for fixation of male-beneficial mutations and
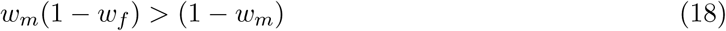

for fixation of female-beneficial alleles. Figure S1 shows the resulting evolutionary dynamics that lead to polymorphism under fitness landscapes with negative curvature. Figure S2 shows an example trajectory for a population evolving under negative curvature. We see that evolutionary trajectories tend towards intermediate expression before gaining polymorphism, just as in Figure 3 of the main text. In the case of negative curvature this results in a decline in population mean fitness as male and female fitness equalize (Fig. S2).

### Individual-based simulations of regulatory evolution

We carried out evolutionary simulations for a single gene and multi-gene cascades using the model for transcription factor binding described in the main text. We simulated populations evolving under the diploid Moran model^44^ [REF Moran 1953a and b] with sexual reproduction. Populations are composed of *N*/2 diploid males and *N*/2 diploid females. The model captures the case of a fixed population size with non-overlapping generations.

The model consists of birth-death events at each time step. Reproduction events occur by choosing one male and one female according to their normalised fitness. These two individuals produce a single offspring who receives one allele from each parent at each locus. The offspring is randomly assigned male or female sex with equal probability and another individual of the same sex is randomly selected (with uniform probability) to die.

Mutation events occur during the transmission of alleles from parents to offspring, with a per nucleotide mutation rate of *µ* = 0. 1/(2*N n*) to ensure weak mutation. We assume that no recombination events occur within TF binding sites, which is justified owing to the short sequence lengths under consideration^30^. In the case of simulated gene cascades, however, recombination can occur between the binding sites of the different genes.

We simulated regulatory evolution under SA selection on the expression of a gene, For cascades, the gene under selection is the terminal gene. We calculated the expression level of each gene in the cascade according to our model of TF binding which then gave the fitness of the individual based on the expression level of the terminal gene. Figure S3 shows the ensemble mean fitness over time for a three-gene cascade as described in the main text. We see dynamics similar to Figure S2 with male and female fitness equalizing at the expense of declining population mean fitness. However we the see a subsequent increase in mean fitness as displacement occurs (see Figure 4 of the main text).

### Displaced polymorphism

In the evolutionary dynamics described in Figure 4 of the main text we see polymorphism arise first at the terminal gene under direct SA selection, before being displaced to regulatory genes higher up the cascade. In particular, there is a tendency for polymorphism to be displaced ultimately to the highest gene in the cascade. This displacement of polymorphism raises three questions. First, why does polymorphism tend to arise at the terminal gene initially? Second, why does polymorphism get displaced? And third, why does polymorphism ultimately rise to the top of the cascade?

The initial emergence of polymorphism at the terminal gene cam be explained by the effect of mutations at the three levels of the cascade on terminal expression levels. We assume initially that all binding sites are functional and genes are highly expressed. As shown in Figure 3 of the main text and Figure S1 above, selection is then essentially directional and favors decreasing expression at the terminal gene. This is most effectively achieved by decreasing binding strength at the terminal gene, since mutations at points higher in the cascade are of smaller effect. Regulatory variants therefore invade most likely at the binding site of the terminal gene and disruptive selection then leads to polymorphism at the bottom of the cascade.

Once regulatory polymorphism is established at the terminal gene, displacement occurs to mitigate the deleterious effects of intermediate expression in heterozygotes. When polymorphism resides at the terminal gene, heterozygotes with one strong and one weak terminal binding site have expression of approximately *E* = 1/2. In the fitness landscape with negative curvature, this is the lowest possible fitness. If polymorphism resides at a point higher in the cascade, in contrast, it is either gene 2 or gene 3 that has expression *E* = 1/2 in the heterozygote. This typically results in expression other than 1/2 in the terminal gene, and therefore greater heterozygote fitness. It is this fitness difference that drives the displacement of polymorphism. Accordingly, there is an increase in mean fitness as polymorphism gets displaced (see Fig. S3 and Fig. 4 of the main text) and cascades with polymorphism residing further up have higher fitness (Figs S4 and S5). Obviously, the advantage of superior heterozygote fitness would also be expected to favour the initial establishment of polymorphism at gene 2 or 3, rather than the terminal gene. But as explained above, this is less likely to occur due to the smaller mutational effects, and hence reduced probabilities of invasion, of regulatory mutations in the bindings sites of genes 2 and 3.

A remarkable result of our simulations is the fact that polymorphism always tends to rise to the top of the cascade and ultimately reside at gene 3 (Fig. 4, main text). At first sight, this is surprising, because there is no mean fitness advantage to polymorphism at gene 3 as compared to gene 2 (see Fig. S4). The difference polymorphisms at gene 2 and gene 3 is related to the resulting sex-specific fitness, as shown in Figures S5 and S6. Here we can see that polymorphism residing at gene 2 typically results in a greater difference between male and female fitness than polymorphism resulting at gene 3. This implies that polymorphism at gene 2 does not totally remove sexual conflict, and as a result there is still disruptive selection that can favour displacement of polymorphism higher up the cascade. Polymorphism at gene 3 then allows for further balancing of male and female fitness, because there is more scope for “fine tuning” expression via gene 2.

The resulting dynamics of polymorphism in a cascade are summarized in Figure S7—there is an inevitable displacement away from the terminal gene as this results in greater heterozygote fitness. There is also a tendency to displace polymorphism to gene 3 rather than gene 2 as this reduces overall sexual conflict via fine tuning of expression lower in the chain.

---

**Figure S1:**
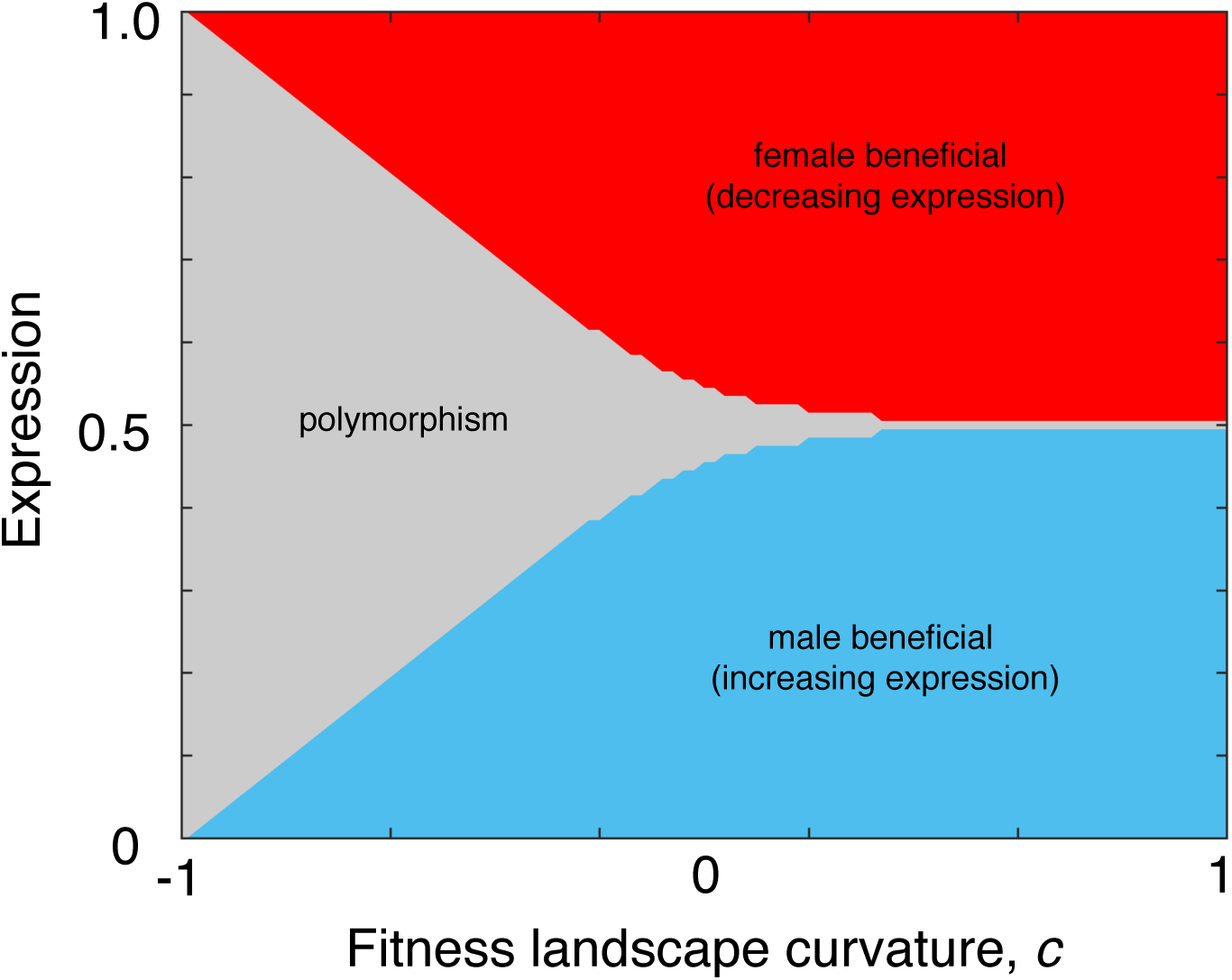
Evolutionary dynamics in the continuum limit. We determined the selection gradient on gene expression using Equations S14 and S15 for the continuum limit under strong selection.We see that expression tends towards intermediate values regardless of landscape curvature, with a large region of disruptive selection leading to polymorphism when curvature is negative.

**Figure S2:**
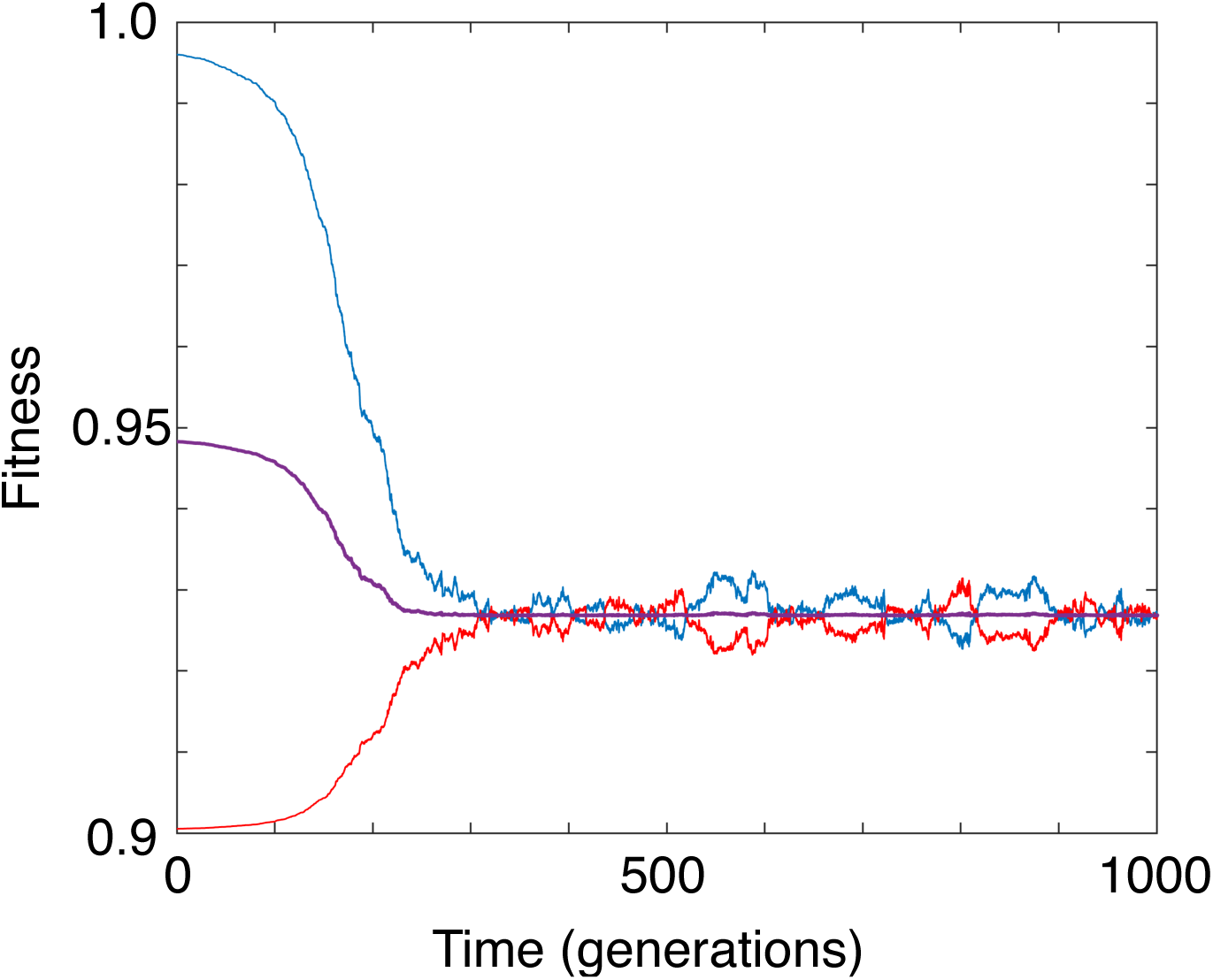
Sample path in the continuum limit. We ran an individual-based simulation in the continuum limit with a population size of N=1000, *s_m_* = *s_f_* = 0.1, *h_m_* = *h_f_* = 10 and *c_m_* = 1 − *c* − *f* = 0.6. We see that mean fitness (purple) declines as male (blue) and female (red) fitness equalize.

**Figure S3:**
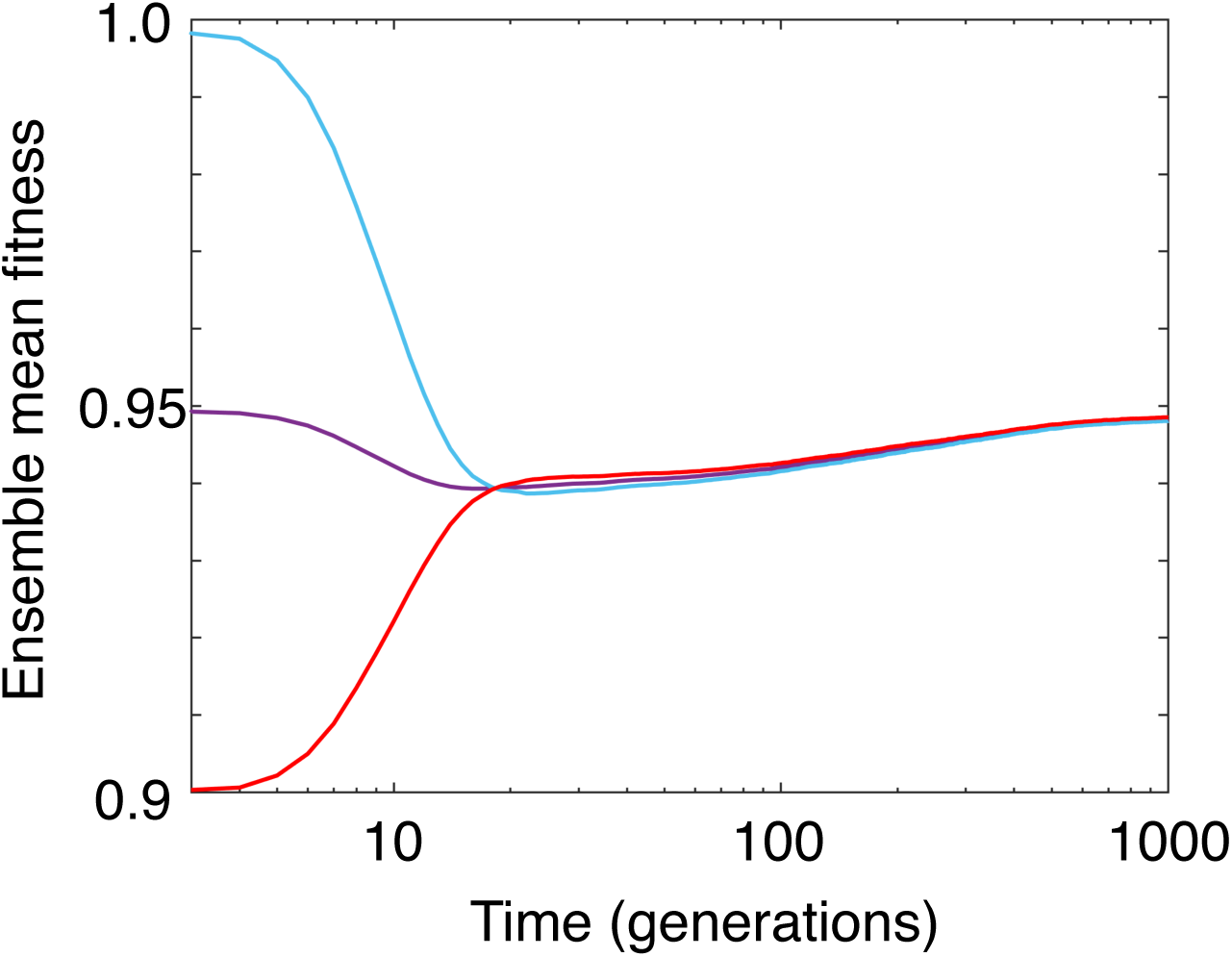
Ensemble mean fitness in a cascade. We plotted the ensemble mean fitness fitness for the whole population (purple), males (blue) and females (red) for the regulatory cascade described in Figure 4 of the main text. We see similar dynamics to Figure S3, with mean fitness declining as male and female fitness equalize. However we also see a subsequent increase in mean fitness and further convergence of male and female fitness as displacement starts to occur.

**Figure S4:**
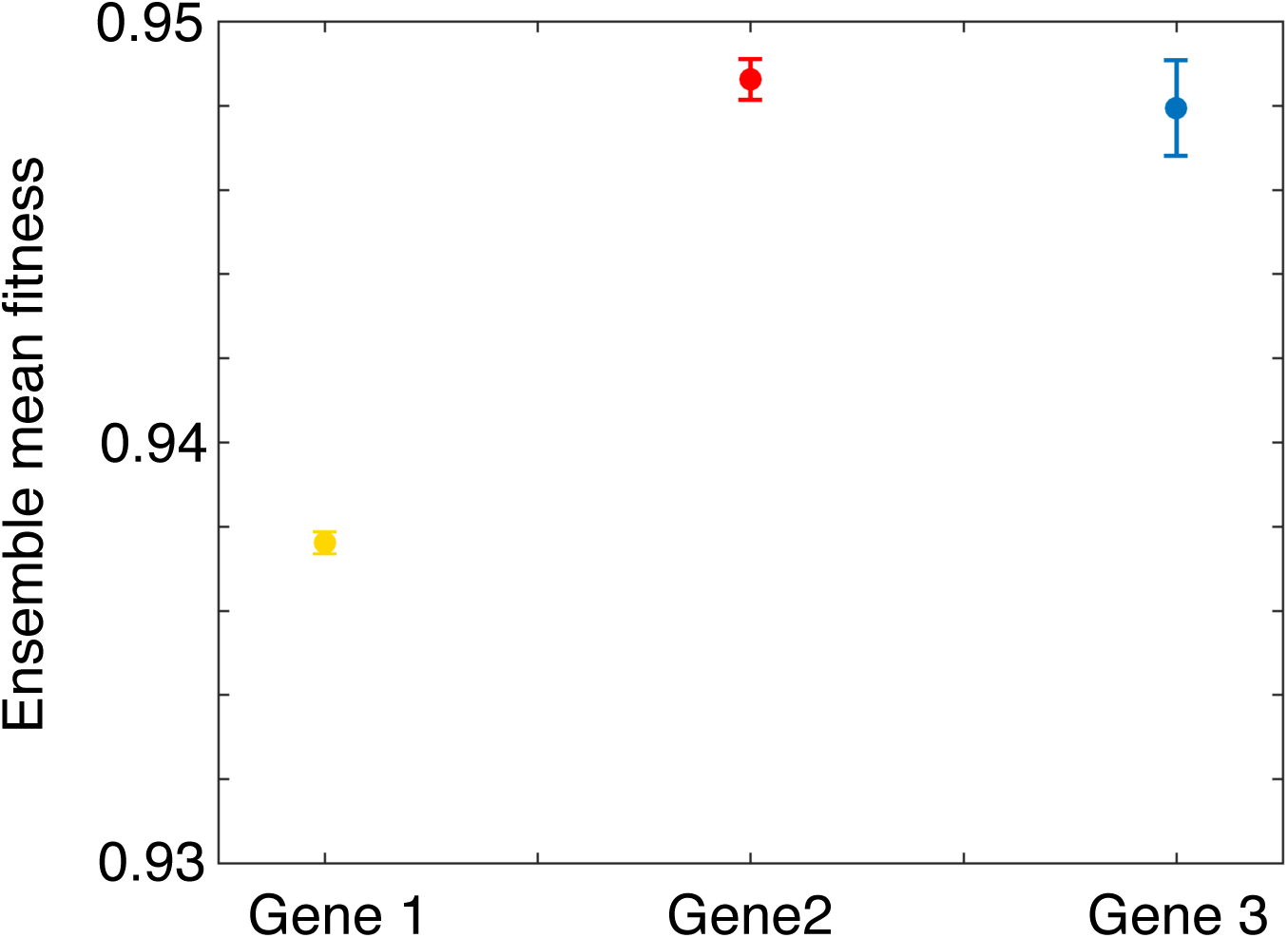
Mean fitness with polymorphisms at different positions in the cascade. We calculated the mean fitness for our cascade simulations for cases where polymorphism resides at the terminal gene, gene 2 or gene 3. We see a clear fitness advantage to polymorphism at genes 2 and 3 compared to the terminal gene 1, but no advantage of polymorphism at gene 3 over polymorphism at gene 2.

**Figure S5:**
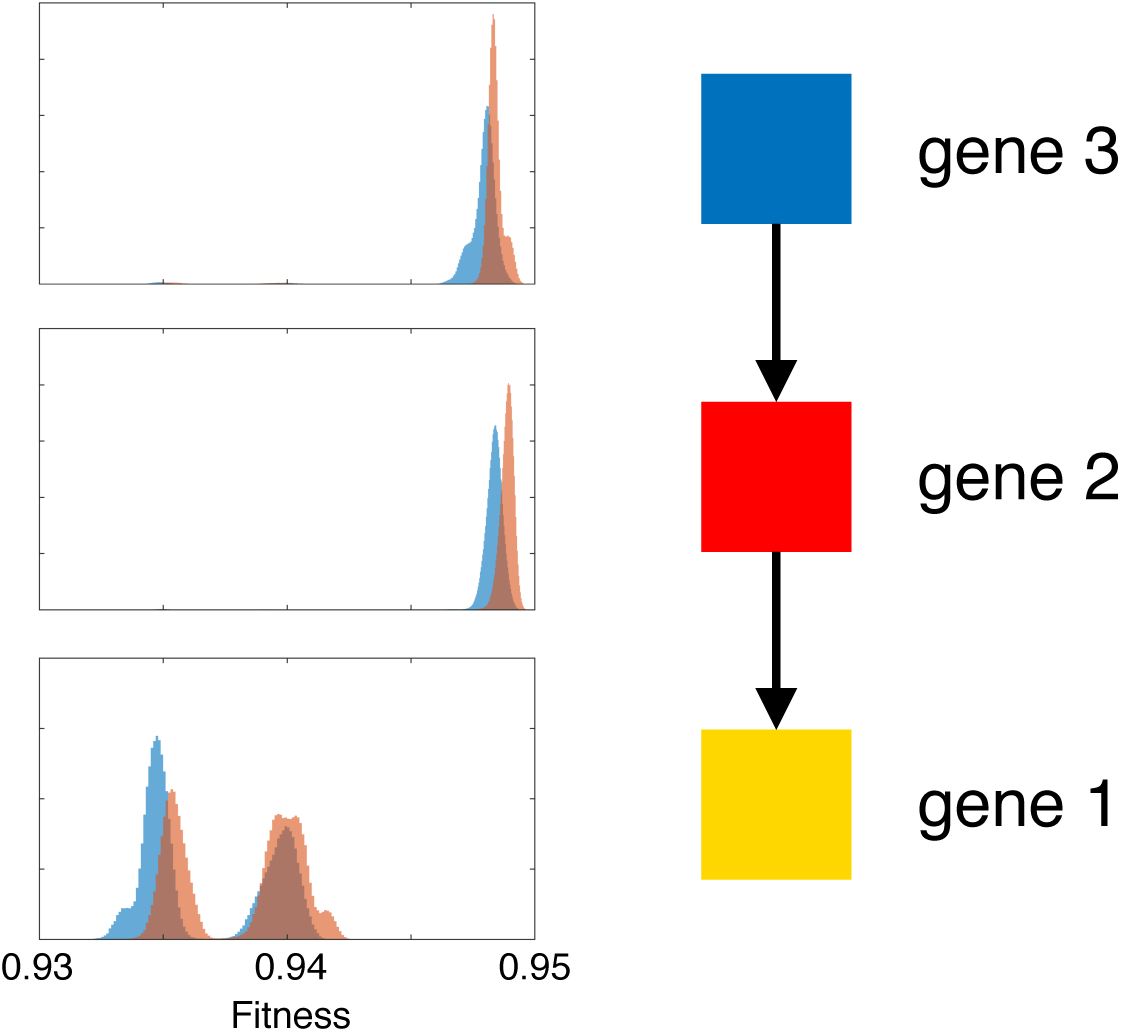
Male and female fitness with polymorphisms at different positions in the cascade. We plotted the distribution of male (blue) and female (red) fitness for populations with polymorphism residing at gene1, gene 2 and gene 3 respectively. We see that gene 1 has both lower average fitness and larger fitness differences between the sexes. Polymorphism at gene 2 results in higher mean fitness and smaller but still prevalent fitness differences between the sexes. Polymorphism at gene 3 results in the lowest fitness difference between the sexes.

**Figure S6:**
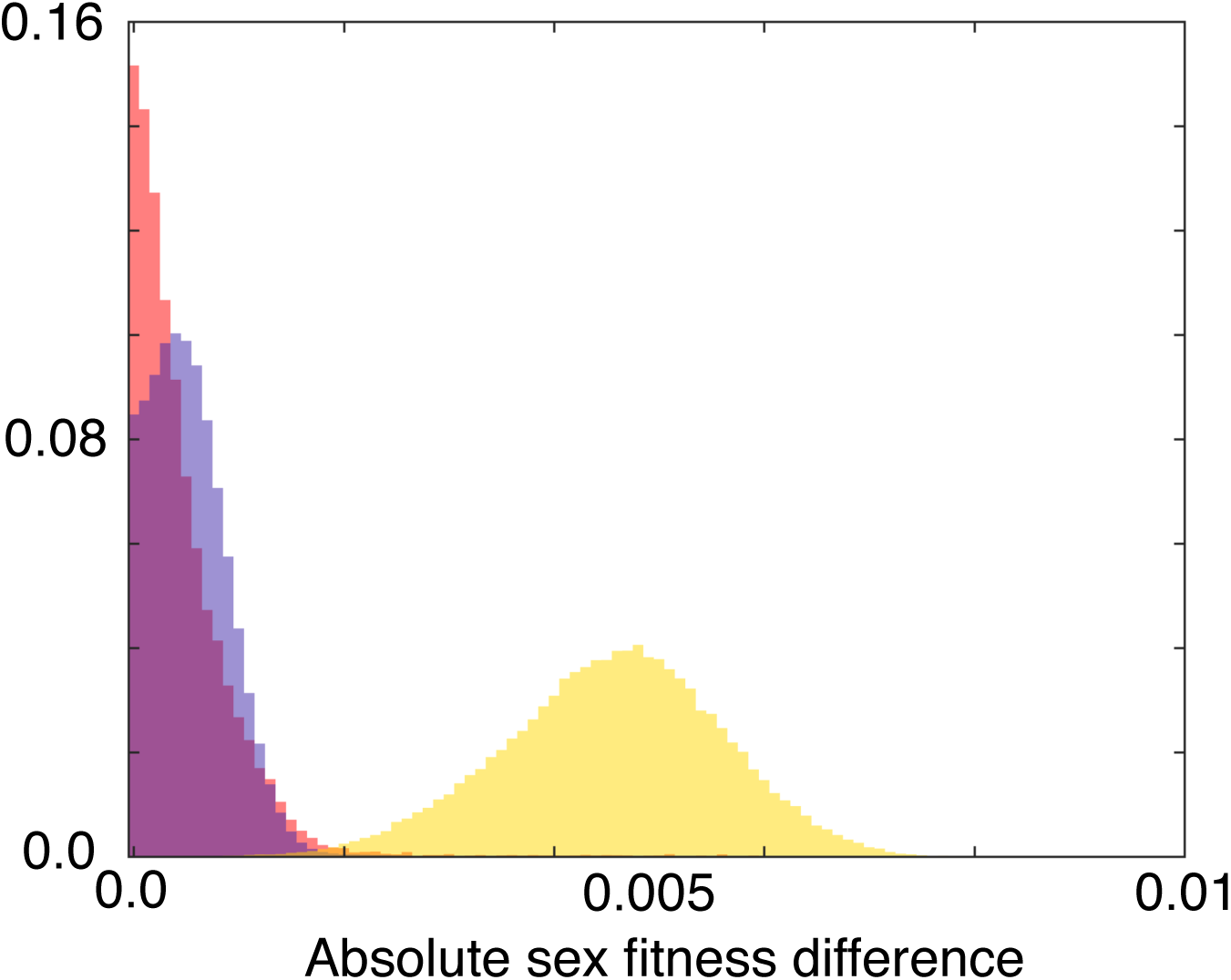
Difference between male and female fitness with polymorphisms at different positions in the cascade. We plotted the absolute fitness difference between males and females for each population with polymorphism residing at gene 1 (yellow), gene 2 (blue) or gene 3 (red). We see that gene 1 always maintains large differences, while gene 2 eliminates differences in some but not most cases, whereas gene 3 the smallest fitness difference between the sexes.

**Figure S7:**
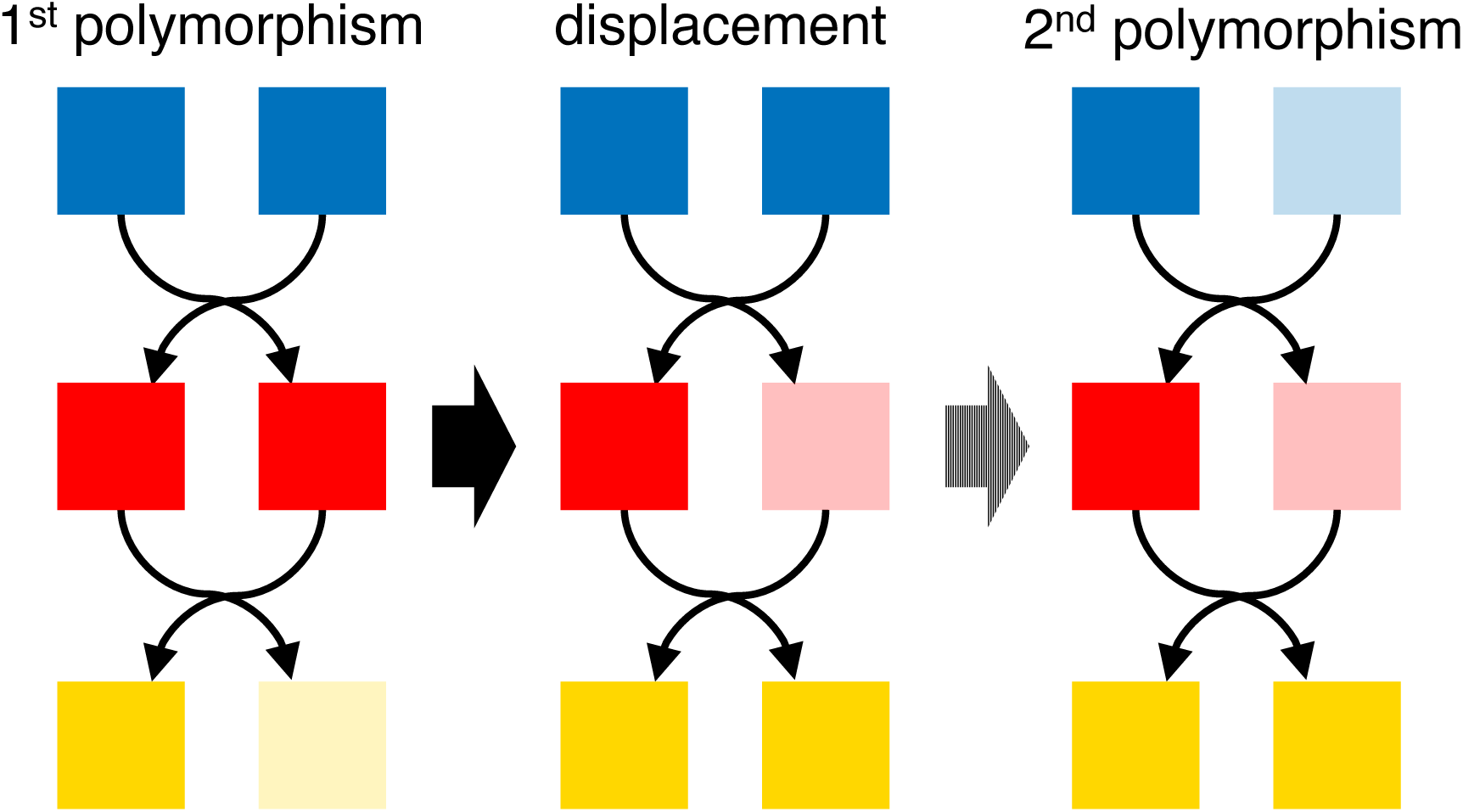
The dynamics of displaced polymorphism. Our study reveals a typical sequence of evolutionary responses to sexually antagonistic selection on gene regulation are as follows. (Left) Polymorphism arises first at the gene under SA selection, where selection is strongest. (Center) polymorphism subsequently gets displaced up the cascade as this delivers a fitness benefit and tends to reduce conflict by equalizing male and female fitness. (Right) When polymorphism is displaced to intermediate levels in the cascade, a second displacement is likely to occur to a point higher in the cascade, as this tends to further reduce conflict.

